# Extreme slowdown of dsDNA and RNA:DNA hybrids in solid-state nanopores for LiCl concentrations above 4 mol/L

**DOI:** 10.64898/2026.10.28.762501

**Authors:** Simon Brauburger, Yunkai Wu, Ulrich F. Keyser

## Abstract

Solid-state nanopores enable single-molecule detection and characterization through ionic-current measurements, but rapid translocation can limit the resolution of molecular features. Here, we investigated the translocation of 7.2 kb dsDNA and 3.3 kb RNA:DNA hybrids at LiCl concentrations extending to 7 mol/L. Both constructs exhibited a substantial increase of ≳ 500% in mean translocation time when increasing LiCl concentration from 2–3 M to 7 M . We also observed a decrease in translocation frequency at increasing LiCl concentration. Both observations indicate a reduction in the effective electric driving force. Further analysis indicates a substantial decrease in the relative width of secondary peaks induced by labels on the molecule. Our results show that increasing salt concentration beyond 4 mol/L is an accessible way for increasing translocation time in solid-state nanopore sensing, with possible implications for barcode sensing resolution.

## 1 Introduction

Solid-state nanopores (SSNPs) provide a platform for single-molecule detection and characterisation, allowing us to investigate sequential and structural features of individual nucleic acids by measuring transient changes in ionic current during their translocation through a voltage-biased pore. The translocation velocity *v*_*t*_ of molecules moving through the pore directly affects the effective resolution. Given the effective sampling limit imposed by high-frequency noise components,^1,2^ a longer translocation time *τ* therefore means more data points per nucleic acid length at a fixed sampling rate. Slower translocation can therefore improve the ability to resolve features along a nucleic acid,^3–7^ which is particularly relevant when the information of interest is encoded in secondary peaks along the molecule.^8^

Using highly concentrated lithium chloride (LiCl) solutions has been established as an effective approach for slowing DNA translocation. Kowalczyk et al. demonstrated a ∼5-fold increase in *τ* when switching from the previous “standard” buffer 1 M KCl to 1 M LiCl (M = mol/L).^9^ Another two-fold increase was obtained when increasing LiCl concentration from 1 to 4 M. Molecular dynamics simulations indicated that this effect was caused by more effective charge neutralization of the DNA. Li^+^ ions fit better into the minor groove, which led to longer-lived binding of Li^+^ ions to DNA, reducing electrophoretic force and resulting in the observed slowdown. However, to the best of our knowledge, the translocation behaviour of DNA in even higher LiCl concentrations has not been characterized yet. Similarly, the impact of changing salt concentrations on the translocation dynamics of RNA:DNA hybrid structures under these extreme conditions remains unexplored.

In this study, we investigate nucleic acid translocation in LiCl solutions at concentrations ranging from 2 M up to 7 M using a 7.2 kb M13 dsDNA and a 3.3 kb RNA:DNA hybrid construct. We found that increasing LiCl concentration from 2–3 M to 7 M increased dwell time by ≳ 500% for both constructs. At the same time, the translocation frequency decreased. For the DNA construct, we also observed a decrease in the fraction of the total translocation time occupied by label-associated secondary peaks. These results extend the available experimental data on concentration-dependent nucleic acid transport and demonstrate the potential of highly concentrated LiCl solutions for further tuning translocation dynamics in nanopore sensing.

## 2 Results

The experimental setup and the two constructs are sketched in Fig. 1a. Linearized M13 ssDNA was hybridised with short complementary DNA oligonucleotides to form dsDNA, containing 6 binding regions with 10 dumbbells each. A 3.3 kb ssRNA fragment was hybridised to form a RNA:DNA hybrid containing 4 binding regions with 6 dumbbells each. A shorter RNA molecule was chosen to represent the usually shorter target lengths of e.g. mRNA. Measurements were performed using multiple SSNPs at an applied voltage of 600 mV and a temperature of 20 ^*°*^C. The samples were measured sequentially. Where possible, multiple measurements with different LiCl concentrations were performed in the same pore (see Methods section).

**Figure 1:**
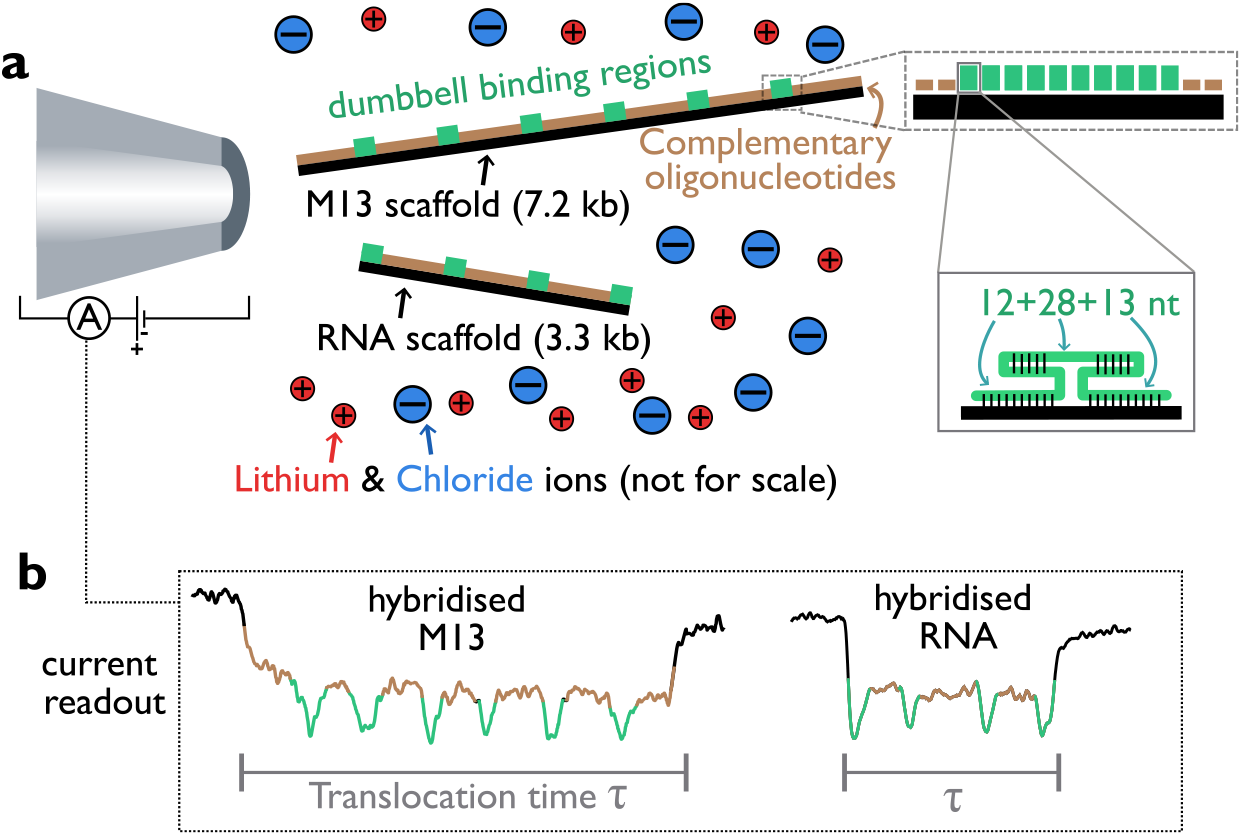
Experimental setup. (a) Schematic of the detection of dsDNA and RNA:DNA hybrid constructs via a solid-state nanopore under an applied electric field. (b) Ionic-current readout of characteristic current blockade signatures during the translocation of hybridized DNA and RNA molecules. The green downward spikes represent localized current modulations associated with the labelled regions.

The dependence of *τ* on LiCl concentration is shown in Fig. 2a+b. Each data point represents the arithmetic mean translocation time (see Fig. 1b) of the events recorded from an individual nanopore at a given concentration. Both constructs exhibited substantial increases in *τ* with increasing LiCl concentration. For the DNA-based construct (Fig. 2a), mean translocation time averaged across pores increased from 937 ± 54 *µ*s at 4 M LiCl (*n* = 3 pores) to 5658 ± 1863 *µ*s at 7 M (*n* = 2 pores). For the RNA-based construct (Fig. 2b), the corresponding values increased from 157 ± 7 *µ*s at 2 M (*n* = 3 pores) to 1004 ± 272 *µ*s at 7 M (*n* = 4 pores). These trends were also observed within individual nanopores. For example, the mean translocation time for hybridized DNA in pore 1 increased from approximately 622 *µ*s at 3 M to 4341 *µ*s at 7 M, corresponding to a 7.00-fold increase. Similarly, the mean translocation time for hybridized RNA in pore 5 increased from approximately 165 *µ*s at 2 M to 777 *µ*s at 5 M, corresponding to a 4.7-fold increase. While the magnitude of the effect appears to depend on the specific pore, our observation of slowdown in different nanopores excludes pore-specific effects as the sole explanation for the observed increase. To illustrate the slowdown, two exemplary DNA events for both 4 M and 7 M in the same pore are shown in Fig. 2c+d.

**Figure 2:**
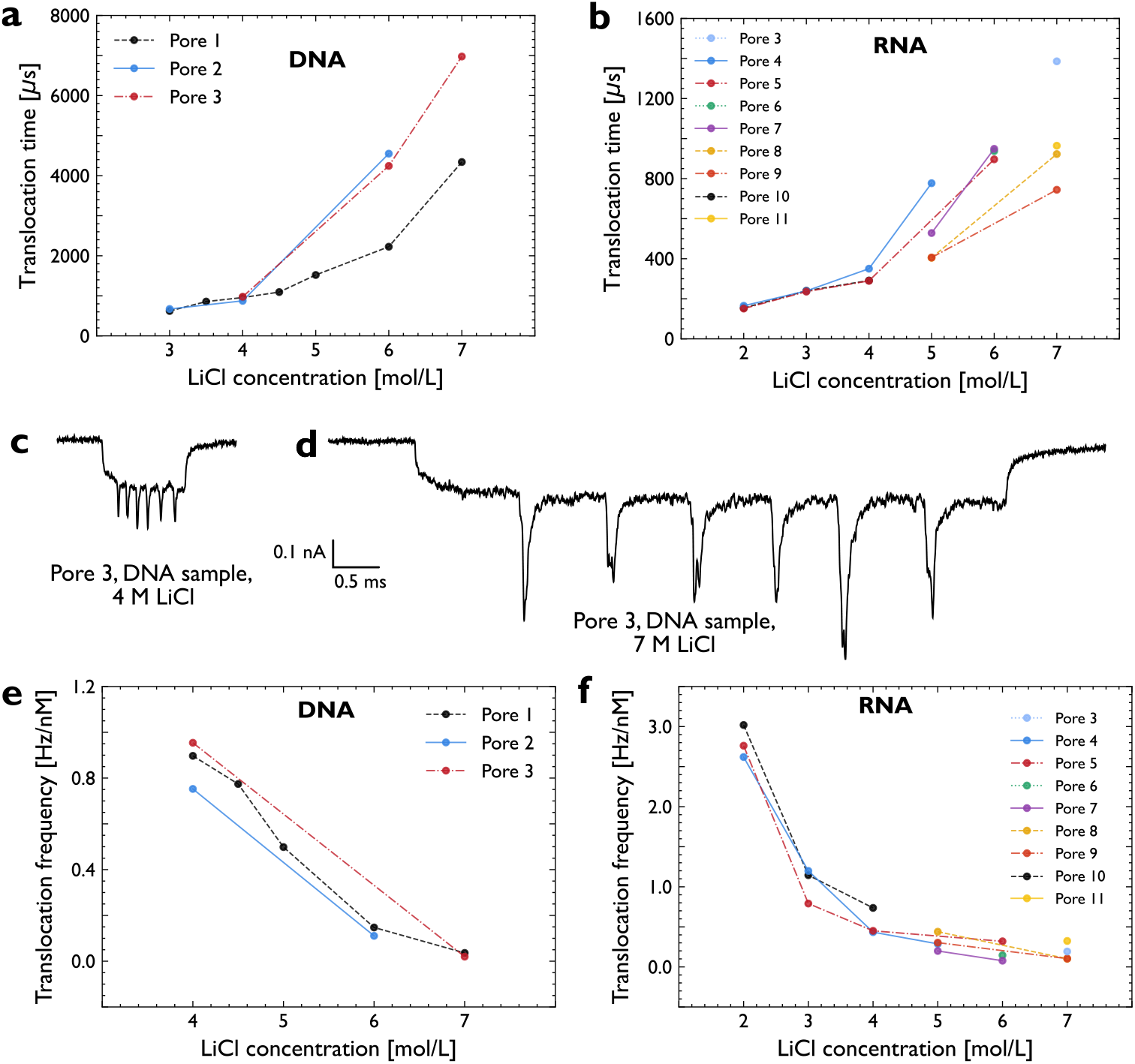
Effect of LiCl concentration on translocation time and frequency. (a,b) Mean translocation times at different LiCl concentrations for the DNA and RNA constructs, respectively. (c) Exemplary unfolded DNA construct event at 4 M LiCl; (d) Exemplary event at 7 M LiCl, recorded using the same sample and pore, displayed on the same scale. (e,f) Mean normalized translocation frequency at different LiCl concentrations for DNA and RNA construct, respectively. Not all measurements that were used for *τ* were suitable for a frequency analysis due to partial clogging or sticking events, and have therefore been excluded in e+f.

At higher LiCl concentrations, relative pore-to-pore variability increased substantially. For the three DNA nanopores measured at both 4 and 6 M, the mean translocation time and standard deviation across pores increased from 937 ± 54 *µ*s to 3673 ± 1262 *µ*s, with the corresponding coefficient of variation increasing nearly sixfold, from 5.8% to 34.4%. At the same time, all pores had relatively similar resistance values, all lying between 71 and 82 MΩ at 4 M LiCl, corresponding to ≈ 7–8 nm following previous model calculations.^10^

In contrast to the increasing translocation time, the translocation frequency decreased at higher LiCl concentrations for both molecule types, as shown in Fig. 2e+f. Measurements were also attempted in 8 M LiCl, but the capture rate at this concentration was too low to give sufficient events for analysis.

The exemplary traces in Fig. 2c+d suggest that the slowdown may affect the properties of the secondary peaks induced by the labelled regions. To test this hypothesis, we investigated the effect of LiCl concentration on peak width in Fig. 3. First, we measured the full widths at half-maximum (FWHM), *t*_*i*_, for the 6 peaks in the DNA sample as shown in Fig. 3a. We then determined the peak length fraction *f* by dividing the sum of *t*_*i*_ by the translocation time of that trace. The median value for each of the DNA measurements is shown in Fig. 3b. For all 3 pores, *f* roughly halves when moving from 4 M to 6 or 7 M.

**Figure 3:**
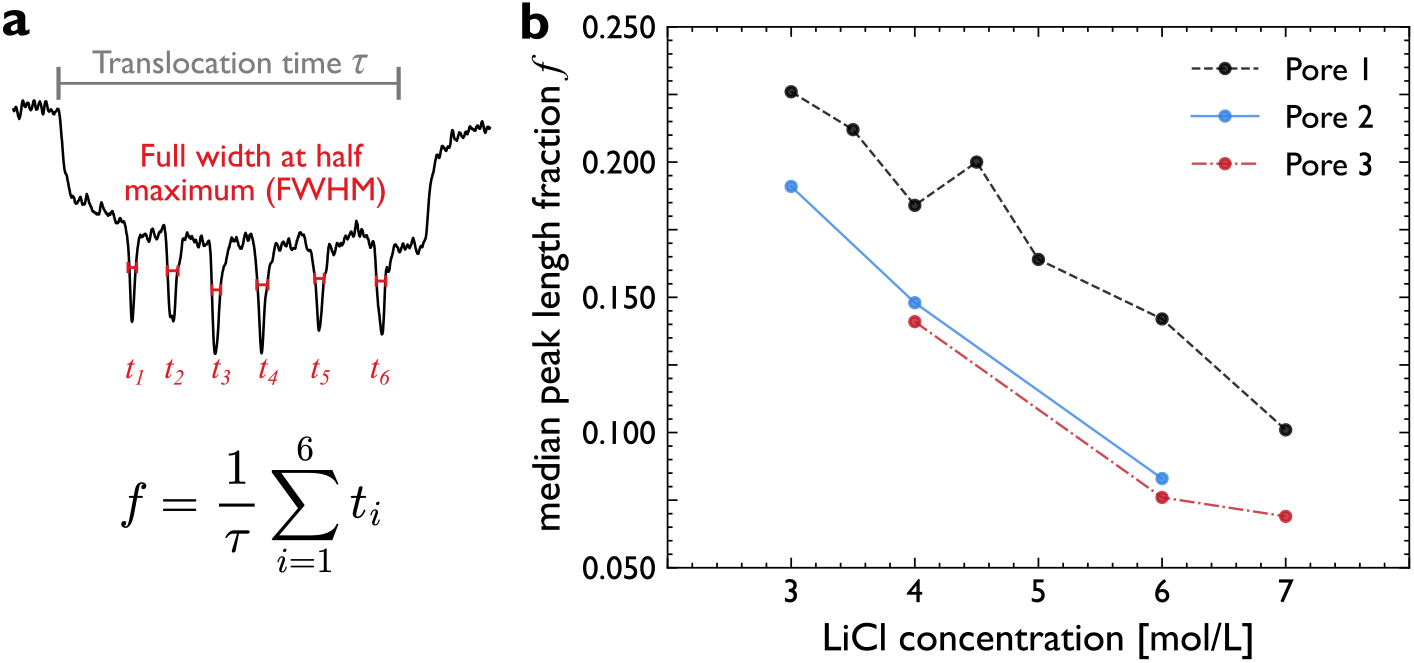
Effect of LiCl concentration on peak width. (a) Definition of peak length fraction *f* and sketch of constituent parts. (b) Change of median *f* with LiCl concentration *c* for the DNA sample. A decrease in *f* with *c* is observed in all three pores. Same data as in Figure 2a+e.

## 3 Discussion

Our results demonstrate a pronounced increase in DNA and RNA:DNA hybrid translocation times at LiCl concentrations above 4 M, extending the concentration range previously investigated by Kowalczyk et al.^9^ This slowdown is accompanied by a substantial decrease in translocation frequency. This inverse relationship between translocation time and frequency follows the trend observed at lower salt concentrations, but also when varying other experimental parameters such as voltage, pressure or pH.^2,11,12^ Our results therefore suggest a further reduction in the effective electrophoretic driving force beyond 4 M LiCl, potentially due to increased counterion screening. The observed increase in pore-to-pore variability at higher concentrations also matches that picture: pore-specific conditions may have a larger relative effect on the driving force if the involved forces are generally of lower magnitude.

The widths of secondary peaks associated with the dumbbell-labelled regions in the DNA sample do not scale in the same way with LiCl as the overall translocation time. Instead, their relative temporal widths compared to total event duration decrease by approximately half between 4 and 7 M LiCl. This suggests that the slowdown may affect different regions of the molecular construct differently, in line with previous observations of local velocity modulations in labelled areas.^2^

From an application perspective, both the general slowdown and the smaller relative widths of the peaks could facilitate unprecedented resolution of labelled nucleic acid targets in SSNPs. For example, the bit density of barcoded molecules^8,13^ could be increased, which is highly relevant for applications such as DNA data storage.^14^ Further experiments and characterization are needed to quantify the possible resolution increase. Higher label density could also expand the range of feasible biological targets, for example short mRNA. It is also important to note that the increased translocation time comes at the expense of reduced capture frequency, introducing a trade-off between temporal resolution and measurement throughput.^2^ While highly concentrated LiCl may therefore be particularly useful for applications requiring extended observation times, the optimal LiCl concentration will depend on the requirements of the measurement.

## 4 Conclusion

We have investigated the translocation of dsDNA and RNA:DNA hybrid constructs through SSNPs in various LiCl concentrations. For both sample types, we demonstrate substantial slowing of translocation time through solid-state nanopores at increasing LiCl concentrations up to 7 M, accompanied by a pronounced reduction in translocation frequency.

Our findings establish this concentration range as a promising avenue for extending molecular observation times and increasing labelling density at the cost of capture efficiency. Our results further motivate investigation of both unexplored parameter regimes and the mechanisms underlying the signal generation process.

## 5 Methods

### 5.1 Scaffold Preparation

The DNA scaffold was prepared following a previously published protocol.^15^ Single-stranded M13mp18 DNA (7249 nt; Guild BioSciences) was linearised using EcoRI-HF and BamHI-HF restriction enzymes (New England Biolabs). Briefly, 8 *µ*L of M13 scaffold at 100 nM was combined with 1 *µ*L of each restriction-site-forming oligonucleotide at 100 *µ*M in 1× CutSmart buffer. The mixture was heated to 70 ^*°*^C for 30 s and cooled to room temperature over 40 min. Following addition of 1 *µ*L of each restriction enzyme, digestion was performed at 37 ^*°*^C for 1 h. The resulting linearised scaffold (7228 nt) was purified using a Monarch PCR and plasmid DNA purification kit and eluted in 10 mM Tris–HCl, pH 8.0. DNA concentration was determined using a NanoDrop 2000 spectrophotometer (Thermo Fisher Scientific).

The RNA scaffold is a copy of the first segment of the Rotavirus RF strain, and was a generous gift by collaborators.

### 5.2 Preparation of the Constructs

The constructs were assembled by hybridising complementary oligonucleotides to the linearised scaffold. For both constructs, dumbbells were used as labels. For DNA, we used a dumbbell-carrier design described previously,^15^ comprising six labelled regions, each containing ten dumb-bell binding sites, spaced 750 bp apart along the scaffold, as sketched in Fig. 1a. For RNA, we used a new design comprising four labelled regions, each comprising six dumbbell binding sites, spaced 680 bp (in between peak 1+2 and 3+4) and 980 bp (between central two peaks 2+3) apart along the RNA scaffold. Oligonucleotides were purchased from Integrated DNA Technologies. Assembly reactions contained 20 nM linearised scaffold and a threefold molar excess of the constituent oligonucleotides (60 nM) in a total volume of 40 *µ*L. The assembly buffer contained 10 mM Tris–HCl, pH 8.0, and 10 mM MgCl_2_. Samples were heated to 70 ^*°*^C for 5 min and cooled linearly to 25 ^*°*^C over 45 min. Excess oligonucleotides were removed using Amicon Ultra 0.5 mL centrifugal filters with a 100 kDa molecular-weight cutoff (Millipore). Samples were washed twice with 10 mM Tris–HCl and 0.5 mM MgCl_2_, pH 8.0. The assembled constructs were recovered by inverting the filters and centrifuging at 1000 × *g* for 2 min. Construct concentrations were determined by NanoDrop spectrophotometry, and samples were stored at 4 ^*°*^C until measurement.

### 5.3 Fabrication of Nanopores

Nanopores were fabricated by laser-assisted pulling (P-2000, Sutter Instrument) of quartz glass capillaries (0.5 mm outer diameter, 0.2 mm inner diameter; Sutter Instrument). The resulting nanopipettes were then integrated into polydimethylsiloxane (PDMS) microfluidic chips using 10:1 base-to-curing agent ratio (Sylgard 184, Dow Corning), plasma-bonded (Femto, Diener Electronic) to glass slides, and filled with different concentrations of LiCl (pH 9). Prior to sample introduction, the nanopores were characterized via current–voltage (*I*–*V* ) measurements from -600 to +600 mV using Ag/AgCl electrodes, yielding an estimated pore diameter of 10±2 nm.

### 5.4 Nanopore Measurements

LiCl measurement solutions (Sigma-Aldrich, ≥ 99% purity) were prepared at the required concentrations and adjusted to pH 9. The same LiCl concentration was maintained on both sides of the nanopore during each measurement. Reported electrolyte concentrations correspond to the final LiCl concentrations after accounting for dilution upon addition of the nucleic acid samples. DNA and RNA constructs were added at final concentrations varying from 0.3 nM to 1.2 nM.

Ionic-current measurements were performed using an Axopatch 200B amplifier (Molecular Devices). Signals were low-pass filtered at 50 kHz using an eight-pole Bessel filter (900CT, Frequency Devices) and digitised at 1 MHz using a National Instruments PCIe-6251 acquisition card. A constant voltage of 600 mV was applied, with the cis electrode held at ground potential. Measurements included in the concentration-dependence analysis were performed at 20 ^*°*^C. The temperature was held constant using the setup described in^16^ to compensate for chip heating, which could distort results. Where multiple concentrations were investigated using the same nanopore, the electrolyte was exchanged as described in the next Subsection 5.5.

### 5.5 Refilling of Chips

To investigate the effect of LiCl concentration within individual nanopores, the same chips were reused for sequential measurements at different electrolyte concentrations. Following each measurement, the electrolyte was removed from all chambers, and each chamber was flushed twice with 50 *µ*L of LiCl solution at the concentration required for the next measurement. The chambers were subsequently filled with the same LiCl solution. The chips were sealed to minimise evaporation and refrigerated at 6 ^*°*^C for at least 16 h to allow equilibration (for many measurements the waiting time was significantly longer, i.e. 2–3 days). We performed the concentration changes in different ways, to make sure the observed trends are not caused by possible concentration gradients.^17^ For example, pore 2 was filled with 4 M on day 1, with 6 M on day 5, and then with 3 M on day 7. In contrast, pore 3 was filled with 6 M on day 1, 7 M on day 5, and 4 M on day 6. Still, both pores show very similar scaling. After each concentration change, a current–voltage (*I*–*V* ) measurement was performed over a voltage range of −600 to +600 mV to assess the nanopore condition. Occasionally, a drifting/fluctuating baseline current was observed when connecting the chip to the electrodes. In those cases, the measurement was delayed for 1–3 minutes to allow the current to stabilize.

### 5.6 Event Selection and Translocation Time Analysis

Translocation events were recorded and analysed using custom LabVIEW and Python software (TraceTamer, unpublished). Only unfolded translocation events were retained for the translocation time analysis. The conventions for determining translocation time (*τ* ) and selecting unfolded events are the same as described in a previous publication.^15^ Events identified as folded translocations, kickouts, fragmented molecules, or otherwise inconsistent with typical translocation characteristics were excluded. Each data point in Figure 2a+b represents the arithmetic mean translocation time of all retained unfolded events for the corresponding nanopore and LiCl concentration.

### 5.7 Translocation Frequency

For the translocation frequency analysis, we included both unfolded and folded translocations, but rejected translocations identified as stuck, fragmented, kickouts, and other spurious signals. Translocation frequencies were determined from the timestamps of accepted events for each measurement, including both unfolded and folded translocations. Events that could not be identified as being caused by a translocating nucleic acid sample were omitted. Measurements with too many kickouts or unstable baselines were excluded from the analysis completely. For the retained events, the intervals between successive accepted events were calculated using Δ*t*_*i*_ = *t*_*i*+1_ − *t*_*i*_. The translocation frequency was then calculated using the reciprocal of the mean inter-event interval, 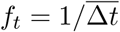. To account for difference between sample concentrations across measurements, frequencies were normalized by the sample concentration and reported in Hz/nM. This normalization is justified by the linear dependence between translocation frequency and sample concentration.^18^

### 5.8 Full Width at Half Maximum Calculation

Follows the protocol described previously^15^ (see supplementary information of the respective publication for extended description). The peak positions and heights are determined via the `scipy.signal.find_peaks` Python function. The estimate of the initial width is then refined by moving outward from the spike minimum in both directions on a smoothed trace and detecting sustained gradient sign changes, obtaining the left/start (green) and right/stop (red) indices for each of the 6 spikes (only for DNA sample). A linear curve is then fit through these 12 points. The “half maximum” absolute value for each spike is then in between the highest absolute y-value of that spike and the absolute y-value of the linear curve at the corresponding x-value. Again moving outward from the peak maximum, the last indices on the right and left side above this absolute value on the side of the maximum are selected. The difference between the right and left indices corresponds to the full width at half maximum.

## 6 Acknowledgments

We thank Alex Borodavka and Cyril Haller for providing the Rotavirus S1 scaffold. U.F.K. and S.B. acknowledge funding through EP/X038009/1 Horizon Europe UKRI Underwrite MSCA ‘DYNAMO’.

